# Assessing RNA as a tool to detect scavenging in a riparian invertebrate predator

**DOI:** 10.64898/2026.08.07.743337

**Authors:** André Rüschendorf, Franziska Middendorf, Jens Schirmel, Bernhard Eitzinger

## Abstract

Riparian environments are characterised by a high diversity of arthropod species, linking the aquatic with the terrestrial ecosystem. One of the dominant arthropod predators in this ecotone, carabid beetles are of particular interest as they may also act as facultative scavengers, feeding on carrion deposited along the shoreline. To test whether carabids feed on carrion, we examined the feeding preferences of the abundant riparian carabid *Bembidion elongatum* in a laboratory feeding trial, offering freshly killed and 24 hours post mortem *Drosophila melanogaster*. We subsequently assessed the detection probability of ribosomal prey RNA and DNA in predator gut contents using *Drosophila*-specific RT-PCR and PCR assays, and quantified nucleotide abundance by quantitative real-time PCR at 0, 3, 6, and 12 hours post-feeding. In the feeding experiment, *B. elongatum* showed a significant preference for fresh over carrion prey. Following consumption, the quantities of prey DNA and RNA in the predator’s gut declined over a 12-hour post-feeding period. However, no differences were detected in prey DNA or RNA quantities between individuals fed fresh prey and those fed carrion. Only immediately after consumption was the DNA:RNA ratio significantly lower in individuals fed fresh prey compared to those fed carrion while this difference was not observed at later time points. Overall, our results indicate that ingested ribosomal prey RNA in predators is present in high quantities, and that the DNA:RNA ratio is not a suitable indicator for distinguishing between consumption of carrion and fresh prey.

## Introduction

Predators in riparian ecosystems link the aquatic with the terrestrial ecosystem by feeding on representatives from both systems (Huszarik et al., 2024; Kripp et al., 2026; Twining et al., 2019). For example, freshwater insects such as chironomid midges emerge from streams in substantial biomass and can contribute significantly to the diets of terrestrial and aerial predators, such as birds, bats and orb-weaving spiders (Middendorf et al., 2025; Schindler & Smits, 2017; Uesugi & Murakami, 2007). While there is wide knowledge on aquatic-terrestrial predator-prey interactions in certain predator groups, such as spiders (Middendorf et al., 2025), much less is known about scavenging, particularly on invertebrate carrion. Invertebrate carrion constitutes an easily accessible resource for numerous arthropod taxa, many of which function as both predators and scavengers (Seastedt et al., 1981). Periodic mass emergences of aquatic insects together with flooding or drought events that deposit dead aquatic invertebrates along shorelines, suggest that invertebrate carrion is likely abundant within riparian ecotones. Consequently, a wide range of consumers is expected to exploit this resource (Orihuela-Torres et al., 2024).

Among arthropod predators, carabid beetles are typically generalist feeders, including carrion in their diet (Lövei & Sunderland, 1996). Studies on riparian carabids indicate that they act both as predators of aquatic and terrestrial prey, and scavengers of carrion originating from aquatic environments that have been washed ashore (Hering & Plachter, 1997). With the exception of a few species specialised in under-water hunting, such as *Carabus variolosus* and *C. clatratus*, carabid beetles are predominantly terrestrial foragers. This suggests that aquatic prey detected in their diet is unlikely to have been captured alive in the water and was more likely consumed after death or upon becoming available on land.

Our knowledge on the feeding ecology of carabid beetles has been substantially enhanced by DNA-based dietary analysis, which allows for the detection and identification of prey remains in field-collected individuals thereby providing insights into otherwise hard-to-observe trophic interactions (Symondson, 2002; Traugott et al., 2013). However, despite its considerable potential, this approach does not permit discrimination between actively captured prey and carrion consumed prior to ingestion (Nielsen et al., 2018). Recent studies suggest that analysing prey RNA may provide a means to differentiate between active predation and scavenging (Cuff et al., 2026; Neidel et al., 2022), potentially extending the benefits of DNA-based dietary analysis in food web research. Because RNA degrades more rapidly than DNA after an organism’s death (Cristescu, 2019), it may allow researchers to distinguish between live and carrion prey at the time of consumption (Neidel et al., 2022).

This study has three aims: First, we want to investigate the food preferences of the riparian carabid beetle *Bembidion elongatum* when offered fresh prey and carrion under experimental laboratory conditions. We expect that *B. elongatum*, similar to other carabid beetles (Hering & Plachter, 1997; Lövei & Sunderland, 1996), will readily accept carrion prey but will prefer fresh prey due to its higher nutritional quality. Second, we want to examine post-consumption detection probabilities of prey RNA and DNA using molecular gut content analysis. We expect that detection probability and quantity of RNA will be higher after consumption of fresh prey than in carrion, and that RNA will exhibit significantly faster temporal decay than DNA. Last, we want to explore the potential application of these findings to distinguish scavenging from active predation in field-based studies. We predict that differences in nucleotide abundance and decay rates will make the DNA:RNA ratio a suitable indicator of carrion feeding.

## Material and methods

### Feeding experiment

Adult riparian carabid beetles (Carabidae: *Bembidion elongatum* DEJEAN 1831, hereafter referred as *Bembidion*) were hand-collected at the Queich river in Landau, Germany (N 49° 12’ 3.2508 E 8° 8’ 3.8436) in October 2022. All beetles were kept in individual glass containers (diameter 4 cm, 4.5 cm high) equipped with moistened tissue paper. Containers were kept in a climate chamber with a 10:14 day-night rhythm and respective temperature change of 20 *°*C and 10 *°*C and humidity of 65%. Beetles were fed ad libitum with cat food (EDEKA, Germany). Prior to start of the experiment, beetles were starved for 72 h in new glass vessels.

For the feeding experiment we used individuals of *Drosophila melanogaster* (hereafter referred to as *Drosophila*) as prey, which were cultured on a cornmeal-yeast diet at the Institute of Environmental Sciences of the RPTU University of Kaiserslautern-Landau. For the feeding experiment, we randomly assigned 60 starved *Bembidion* beetles to one of three feeding treatments:

In Treatment 1, each of the 20 beetles was provided with two *Drosophila* that had been freeze-killed just prior to the start of the experiment. Freshly freeze-killed flies (hereafter referred to *fresh prey*) were used to simulate live prey, as using live prey would have made it difficult to standardize feeding times across treatments.

In Treatment 2, each of the 20 beetles received two *Drosophila* flies that had been left decaying for 24 hours at 20 °C, simulating dead carrion prey (hereafter referred to *carrion*).

In Treatment 3, each of the 20 beetles was offered one freshly freeze-killed *Drosophila* and one that had been left decaying for 24 hours at 20 °C. To distinguish which prey had been fed upon, we marked the fresh *Drosophila* in half of the experiments and the decayed *Drosophila* in the other half with a dot using a black permanent marker (Edding, Ahrensburg, Germany). A Fisher’s exact test showed no significant association between the marking and the choice of the specific prey type (*p* = 0.58).

Feeding by *Bembidion* was observed until end of feeding process, which lasted on average 50 minutes. For each beetle, the number of prey individuals consumed (eaten completely or only partially) and the time until the start of feeding was recorded. After feeding had ended, five beetles from each of the three treatments were assigned to one of four subgroups. These subgroups differed in the time elapsed between prey consumption and freezing of the beetles at –80 °C: immediately (timepoint 0), after 3 hours, after 6 hours, or after 12 hours.

### DNA and RNA extraction

To extract both, DNA and RNA from carabid individuals, including their gut contents, we followed the HighGI RNA extraction protocol by Yoshino et al. (2020). Whole frozen specimens were homogenized using sterile pestles and suspended in 200 µL of lysis buffer. The homogenates were centrifuged at 12,000 × g for 2 minutes, and 150 µL of the resulting supernatant was transferred to a new 1.5 mL microcentrifuge tube. For DNA-extraction we used 75 µL of this lysate, which was mixed with 270 µL of AMPure XP magnetic beads (Beckman Coulter, Indianapolis, USA) and 270 µL Isopropanol and incubated for 5 minutes. The magnetic beads, binding both RNA and DNA, were collected using a magnetic stand and the supernatant was discarded. The beads were then washed three times with 800 µL of 85% ethanol, air-dried for 5 minutes, and nucleic acids were eluted in 100 µL of nuclease-free water.

To prevent interference with RNA detection during reverse transcription PCR, we aimed to obtain RNA extracts free of genomic DNA contamination. RNA was isolated using the remaining 75 µL of lysate, following the same protocol as previously described. The only modification was the inclusion of a DNase treatment (DNA-free Kit, rigorous DNase treatment; Thermo Fisher, Waltham, USA) to remove DNA from the magnetic beads prior to the ethanol wash steps. Successful DNA removal in eluted samples was confirmed by the absence of PCR products in subsequent screening reactions. If RNA sample still tested positive for DNA, DNAse treatment was repeated until there was no further detection of amplifiable DNA.

In addition, to test for potential carry-over contamination, each extraction batch (5–13 samples) included one negative control in which water was used instead of tissue.

### Diagnostic PCR

All *Bembidion* samples were tested in screening PCRs for the presence of *Drosophila* prey using genus-specific primers Droso-S391/ Droso-A381, (5′-AAATAACAATACAGGACTCATATCC-3′ and 5′-GTAATACGCTTACATACATAAAGZTAZA-3′; Wolf et al., 2018) targeting a 240 bp fragment of the nuclear 18s rDNA.

For DNA samples, the PCR was conducted in a 10 μL volume, containing 5 μL MyTaq Red Mix PCR mastermix (Bioline, London, UK), 0.5 μM of each primer, 1 μL molecular grade water, and 3 μL of DNA extract. PCR cycling conditions followed protocol by Wolf et al. (2018) with 15 min at 95 °C, 35 cycles of 30 s at 94 °C, 90 s at 62 °C and 60 s at 72 °C, followed by a final elongation at 72 °C for 10 min.

RNA samples were reverse-transcribed (RT) into stable complementary DNA (cDNA) prior to PCR, using the Qiagen OneStep RT-PCR Kit, following the manufacturers protocol. Each 10 μL PCR contained 0.4 μL RT-PCR Enzyme Mix, 2 μL RT-PCR buffer, 0.4 μL dNTP mix, 0.6 μM of each primer, 4 μL RNAse free water and 2 μL of RNA extract. The thermocycling protocol started with a reverse transcription step at 50 °C for 30 min, followed by denaturation at 95 °C for 15 min, 35 cycles of 94 °C for 30 s, 62 °C for 60 s and 72 °C for 60 s, and a final elongation at 72 °C for 10 min.

PCR products were separated using gel electrophoresis using HDGreen (Intas, Göttingen, Germany)-stained agarose gel. All PCR reactions were performed in duplicates. In cases where the two replicates yielded discrepant results, a third PCR followed by gel electrophoresis was conducted.

### Quantitative Real time PCR (qPCR)

To quantify DNA content in RNA and DNA extracts, all carabid samples were additionally subjected to quantitative real-time PCR (qPCR) using the same primers Droso-S391/ Droso-A381. Prior to qPCR, RNA samples were reverse-transcripted (RT-PCR) to cDNA using the Bioline SensiFAST cDNA Synthesis Kit (Meridion Bioscience, Memphis, USA) following the manufacturers protocol: Each 20 µL RT-PCR contained, 4 µL TransAmp buffer, 1 μL Reverse Transcriptase, 10 μL RNAse free water and 5 μL RNA extract. The RT-PCR cycling conditions were: an initial incubation at 25°C for 10 minutes, followed by 42°C for 15 minutes, and a final step at 85°C for 5 minutes.

Quantitative PCR of all cDNA and DNA samples was then conducted in a qPCR cycler (Realplex 4, Software Realplex Version 2.2.0.84, Eppendorf, Germany). Each 10 µL qPCR consisted of 5 μL SYBR-Green (PowerUp SYBR Green Master Mix; Thermo Fisher, Waltham, USA), 0.75 μL of each primer, 1.5 μL molecular grade water, and 2 μL DNA/cDNA extract.

QPCR conditions were 50 °C for 2 min, 95 °C for 2 min, 55 cycles of 95 °C for 15 s, 62 °C for 15 s and 72 °C for 60 s, and a final elongation at 95 °C for 15 s, 60 °C for 15 s, a linear, 20 min temperature increase up to 95°C, and 95°C for 15 s.

To assess the DNA content of cDNA and DNA samples, a series of standard dilutions (10^−4^, 10^C^, 10^−^8, and 10^−10^) of quantified *Drosophila* PCR products (121 µL/ml) were run alongside in the experimental samples in the same qPCR run.

To generate these standards, DNA was extracted from a single *Drosophila* individual using the HighGI RNA extraction protocol described by Yoshino et al. (2020). The extracted DNA was then amplified using the same PCR protocol outlined above, with primers Droso-S391 and Droso-A381.The *Drosophila* PCR products were purified using AMPure XP magnetic beads (Beckman Coulter, Indianapolis, USA) at a ratio of 1:1.8, following the same washing protocol applied to DNA extracts.

The DNA concentration of the purified standard was then quantified using a Qubit 1.0 Fluorometer (Thermo Fisher Scientific, Waltham, MA, USA). Each sample, along with the negative control and the four standard dilutions, was analysed in triplicate.

All data on RNA and DNA detection results can be found in the Supplementary Material Table S1.

### Statistical analysis

The statistical analysis was performed using R Statistical Software (R Core Team, 2025). Data from the mixed prey group experiment were assigned to the fresh prey group when only fresh prey has been consumed and to the carrion prey group when only carrion was ingested. In the feeding experiment, preference for fresh prey or carrion was tested using an exact binomial test.

We modelled the detection probability of prey RNA and prey DNA, (as determined by diagnostic PCR, where consumption was coded as 1 and non-consumption as 0) as functions of time since feeding, prey type (fresh vs. carrion) and their interactions using generalized linear models (GLM) with a binomial distribution. We then calculated the relationship between prey DNA quantity and prey RNA quantity (as quantified by qPCR) with time since feeding, prey type and their interactions using GLMs with a Gaussian distribution. Finally, we modelled the DNA:RNA quantity ratio as a function of time since feeding, prey type and their interaction using a GLM with a Gaussian distribution. Model significance was assessed using likelihood ratio χ^2^ tests.

## Results

In the single prey trials, *Bembidion* consumed on average one fresh *Drosophila* individual (± 0.54 SD) and a maximum of two individuals. For carrion prey, there was an average consumption of 0.9 *Drosophila* (± 0.34 SD) and a maximum of 1.5 prey individuals. In the mixed prey trial, *Bembidion* showed a significantly higher preference for fresh prey (two-sided: *p*= 0.012; one-sided: *p*=0.006) and chose fresh prey in 16 out of 20 cases (80%).

Using diagnostic PCR, prey DNA was successfully amplified in 54 (90%) and prey RNA in 59 *Bembidion* (98%), respectively. The high detection rates of RNA resulted in an extremely unbalanced binary response with only a single negative observation. Because this data structure led to (quasi-) complete separation and unstable parameter estimates in a binomial generalized linear model, no formal statistical comparison among time points or prey types was performed.

The detection rates of DNA decreased over time, although not significantly (χ^2^ = 0.397, p = 0.529). Moreover, prey type did not significantly affect DNA detection rates (χ^2^ =0.056 p = 0.813; Fig. 1). The amount of prey nucleotide quantity, as measured by qPCR ranged from 4.53×10^−5^ ng/µL to 729.67 ng/µL for DNA and from 5.52×10^−4^ ng/µL to 5836.67 ng/µL for RNA, respectively. Prey DNA and RNA quantities declined over time; however, the decrease was statistically significant only for prey DNA (estimate = −1.162 × 10^−5^, p = 0.044), whereas prey RNA showed no significant temporal change (estimate = −2.929 × 10^−7^, p = 0.994; Fig. 2). There was no significant effect of prey type on DNA and RNA concentrations, although estimates suggested lower DNA quantities in fresh prey than in carrion (estimate = −7.611 × 10^−5^, p = 0.133), and higher RNA quantities in fresh prey than in carrion (estimate = 5.324 × 10^−4^, p = 0.163). There was also no significant interaction between time and prey type for both DNA and RNA.

**Figure 1.**
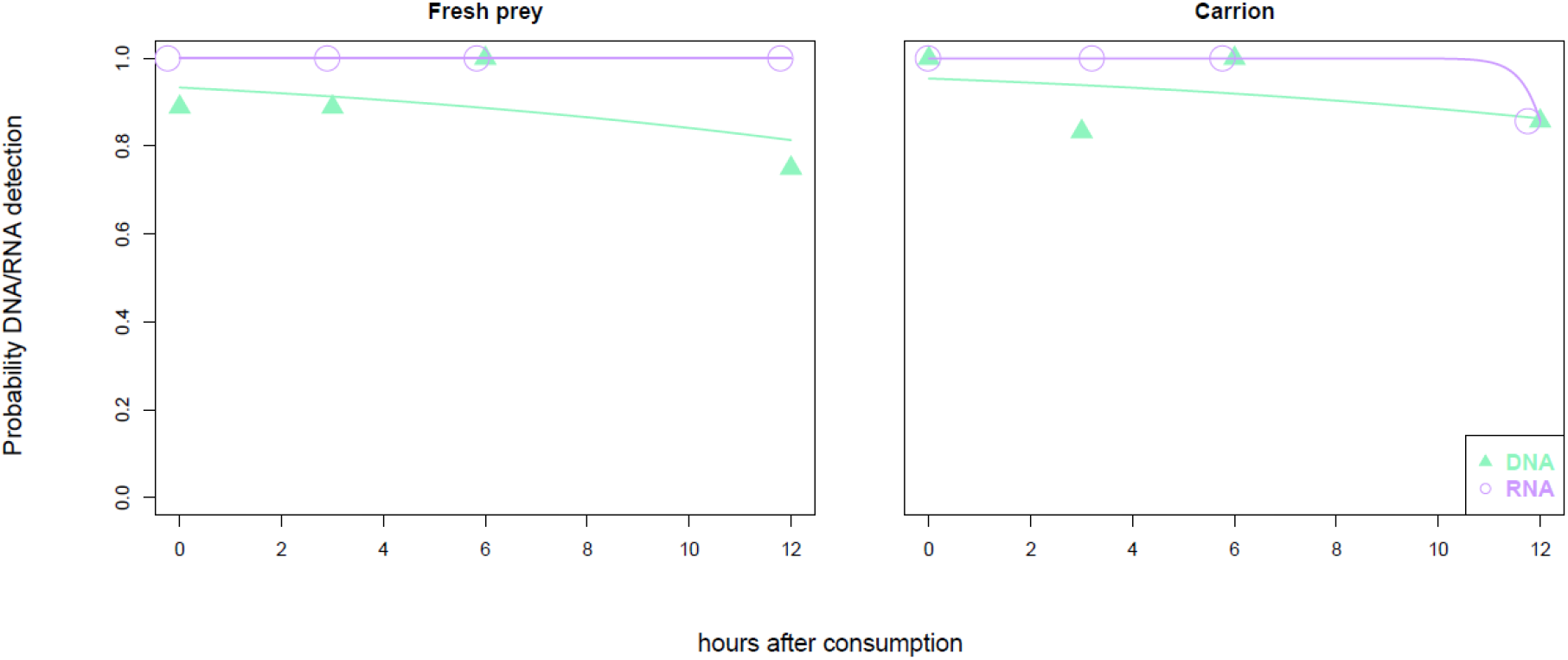
Detection success of prey DNA (green colour) and prey RNA (purple colour) for fresh prey (left panel) and carrion (right panel) from 0 to 12 hours after consumption

**Figure 2.**
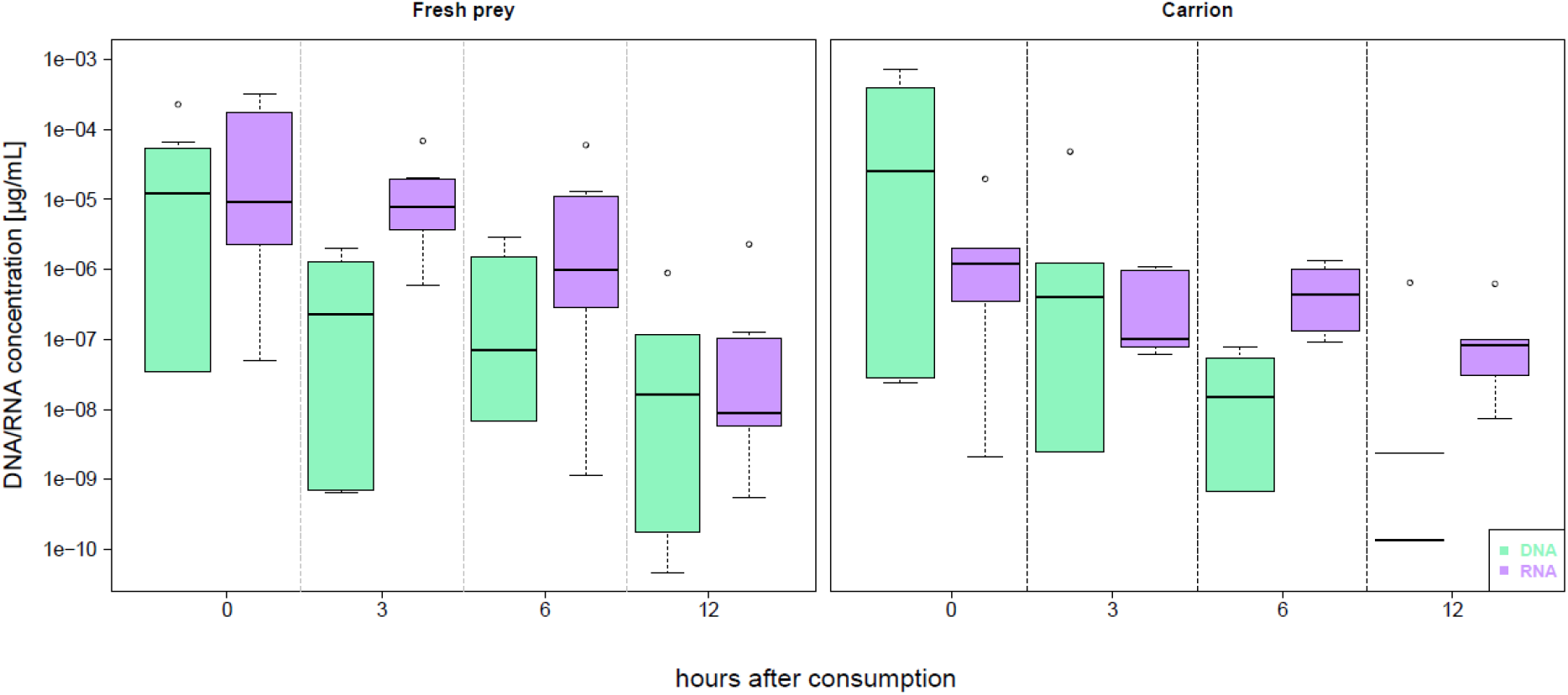
Pairwise comparison of nucleotide quantity (µg/ml) of DNA (green colour) and RNA (purple colour) in fresh prey (left panel) and carrion (right panel) from 0 to 12 hours after consumption.

The DNA:RNA ratio was significantly lower in fresh prey compared to carrion (estimate = –10.97, p = 0.008, Fig. 3). The DNA:RNA ratio decreased significantly over time in carrion (estimate = −1.137, p = 0.013), whereas fresh prey showed a non-significant positive trend (estimate = 0.206, p= 0.572). The significant time × prey type interaction (estimate= 1.343, p = 0.023) indicates that temporal patterns differed between prey types.

**Figure 3.**
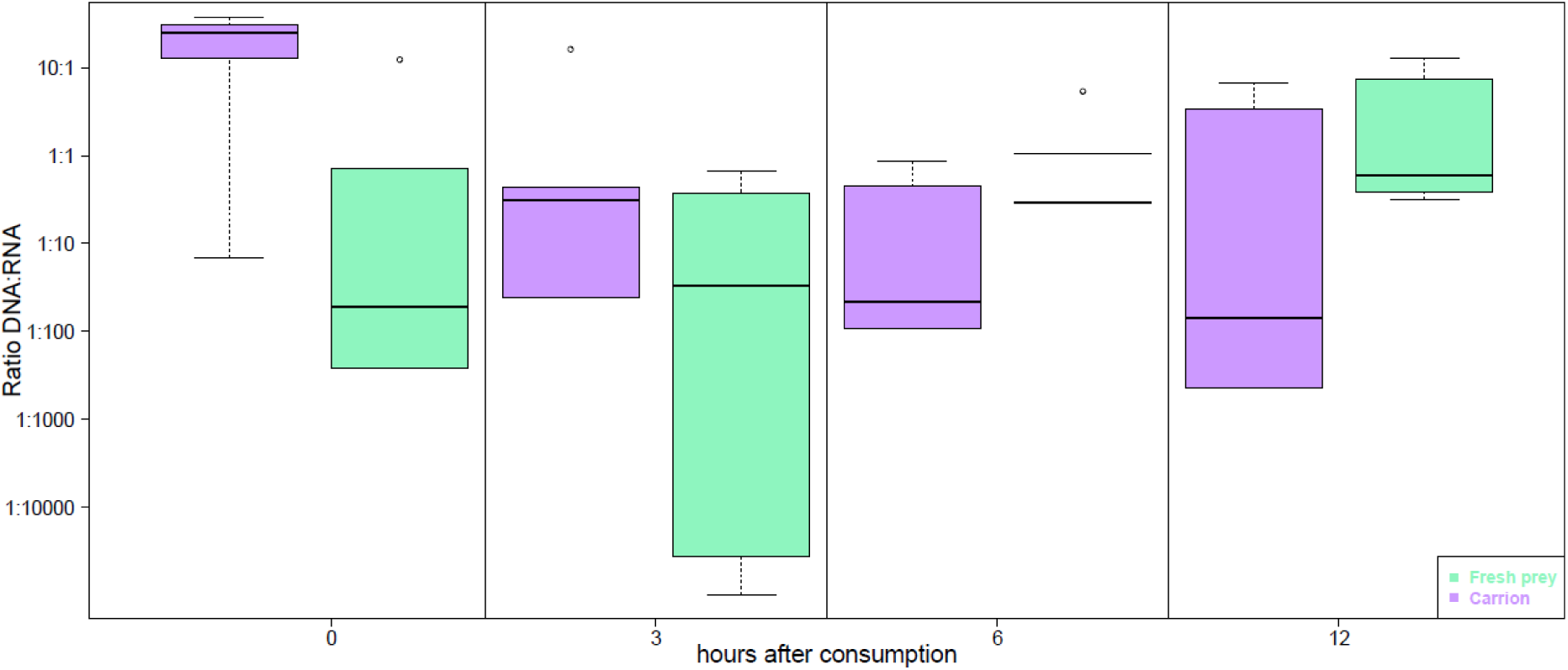
Pairwise comparison in ratio of DNA:RNA of fresh prey (green) and carrion (purple) based on concentration of DNA and RNA from 0 to 12 hours after consumption.

## Discussion

The present study contributes to the growing use of RNA detection in food web studies, that aim to distinguish between active predation on live prey and consumption of carrion by invertebrate predators. Here we found that riparian carabid predator *Bembidion elongatum* accepts both, fresh and carrion prey and that feeding on both can be tracked up to 12 hours after consumption by amplifying RNA and DNA of prey *Drosophila melanogaster*. Furthermore, we found that DNA:RNA ratio to differ between fresh prey and carrion and changed differently over time, indicating that prey condition influences the temporal decay of RNA and DNA.

### Bembidion prefer fresh prey

Our single-prey feeding experiment demonstrates that riparian *Bembidion* beetles readily consume both fresh and carrion prey, indicating that they can be considered facultative scavengers. However, when offered both prey types, beetles consistently preferred fresh prey, confirming our first expectation. This preference may reflect the higher nutritional quality of fresh prey and the progressive loss of nutritional value during decomposition (Foltan et al., 2005). In addition, desiccation may make decomposing prey more difficult to process. Nevertheless, the carrion prey used here was only 24 hours old and therefore represented an early stage of decomposition.

The preference for fresh prey does not preclude scavenging under natural conditions. Riparian carabids frequently consume aquatic invertebrates washed ashore, including chironomids (Hering & Plachter, 1997), and DNA metabarcoding has revealed a diverse diet containing both terrestrial and aquatic prey (Middendorf et al., in revision). Whether fresh prey or carrion is preferred is likely to depend on prey availability, energetic costs of searching and handling, and the nutritional value of the resource. Thus, scavenging may represent an opportunistic feeding strategy that becomes more important when live prey are scarce or carrion is readily available.

### Prey DNA and RNA persist during digestion

We did not find RNA detection rates to be significantly higher in fresh prey than in carrion and RNA quantities were not significantly higher in fresh prey than in carrion. This pattern was consistent across the 12-hour long digestion time and therefore did not support our expectation that RNA would be more rapidly degraded in carrion. These findings contrast with (Neidel et al., 2022) who observed significantly lower RNA detection in predators fed carrion and a rapid decline in RNA detection over time. The difference between the two studies may partly result from the relatively short digestion period used here. We detected prey DNA and RNA in more than 90% of samples throughout the 12-h experiment, indicating that both markers persisted considerably longer than expected. Previous studies have reported prey DNA detection half-lives of approximately 22–34 h in carabid predators (Waldner et al., 2013), while prey DNA can remain detectable for substantially longer periods under some conditions (Neidel & Traugott, 2023). Although we expected the smaller body size and presumably higher metabolic rate of *Bembidion* to result in faster digestion (Ehnes et al., 2011), our results suggest that a 12-h observation period was insufficient to capture substantial differences in RNA persistence between prey types.

The molecular marker used may provide an additional explanation. We targeted a 240-bp fragment of the nuclear 18S rRNA gene, which is highly abundant because ribosomal RNA occurs in large numbers within cells. Consequently, the persistence of 18S rRNA in gut contents may not reflect the persistence of cellular mRNA or other RNA molecules that degrade more rapidly after prey death (Cristescu, 2019). The high abundance of 18S rRNA may therefore obscure differences in RNA degradation between fresh prey and carrion. Similar difficulties have recently been reported by (Junk et al., 2025), who found that 18S rRNA abundance did not allow reliable discrimination between dead food organisms and living endosymbionts in mussels.

Our qPCR results nevertheless provide evidence that DNA and RNA do not necessarily behave identically during digestion. Both DNA and RNA quantities tended to decline over time, but only the decline in DNA was statistically significant. DNA quantities also tended to be higher in carrion, whereas RNA quantities tended to be higher in fresh prey, although neither difference was statistically significant. These results suggest that quantitative measurements of prey nucleic acids may capture temporal changes that are not apparent from simple presence/absence data. However, substantially longer digestion experiments and larger sample sizes will be required to determine whether these trends are reproducible and biologically meaningful.

The DNA:RNA ratio provided a more differentiated picture than either marker alone. We found a significant difference in the DNA:RNA ratio between fresh prey and carrion, with lower ratios in fresh prey. However, this difference was primarily driven by the measurements immediately after feeding and was not maintained at subsequent time points when treatments were examined separately. Furthermore, the ratio decreased significantly over time in carrion, whereas fresh prey showed a positive but non-significant temporal trend. Thus, the significant difference between prey types did not translate into a consistent temporal signature that could reliably identify carrion consumption.

This result differs from (Neidel et al., 2022), who found that RNA:DNA signal ratios differed between fresh prey and carrion and suggested that these ratios could be used to infer scavenging. An important methodological difference is that (Neidel et al., 2022) quantified relative fluorescence units (RFU) obtained through capillary electrophoresis, whereas we quantified DNA and RNA concentrations using qPCR. RFU values provide a semi-quantitative measure of amplicon signal strength and can be influenced by factors such as assay efficiency and appropriate calibration (Thalinger et al., 2021). In contrast, qPCR allows more direct quantitative comparisons of target nucleic acid concentrations. Consequently, the two approaches may not be directly comparable.

Our results therefore suggest that a DNA:RNA ratio based on absolute nucleic acid quantities should not currently be considered a reliable indicator of scavenging. The significant difference between treatments demonstrates that prey condition can influence the relative abundance of DNA and RNA, but the lack of a consistent temporal separation indicates that this signal is insufficiently robust for identifying feeding mode in individual predators. Importantly, the ratio may still have potential under different experimental conditions, particularly with longer digestion periods or alternative molecular markers.

### Is RNA suitable for carrion detection?

Our findings indicate that RNA-based approaches require careful validation before they can be used to distinguish scavenging from predation. The experimental design may have limited our ability to detect differences between prey states because we used freshly killed *Drosophila* to represent fresh prey rather than genuinely live prey. Although this approach allowed feeding and digestion to be standardized, freshly killed prey may retain molecular characteristics very similar to living prey. Likewise, carrion was only 24 h old, potentially limiting differences in initial DNA and RNA content between treatments. Future experiments should therefore include live prey and carrion representing several stages of decomposition to better capture the molecular differences associated with predation and scavenging.

Extending the digestion period would also be important. Our 12-h experiment was chosen based on the assumption that the small body size of *Bembidion* would result in rapid digestion, but the high persistence of both DNA and RNA suggests that longer time intervals are necessary. Longer experiments would allow the relative degradation rates of DNA and RNA to be characterized more accurately and could reveal differences between prey types that were not apparent within the first 12 hours. Such experiments would also better reflect natural conditions, where carabids may experience prolonged periods between feeding events and potentially retain detectable traces of previous meals (Bilde & Toft, 1998).

The choice of molecular marker may be equally important. The 18S locus is highly useful for dietary and biodiversity studies because of its broad taxonomic coverage, conserved primer-binding regions, and high copy number (Albaina et al., 2016; King et al., 2008; Pompanon et al., 2012). However, its high abundance in RNA may make it unsuitable for distinguishing living from dead prey. COI represents a promising alternative because the difference in RNA and DNA copy numbers is substantially smaller than for 18S (Marshall et al., 2021), while extensive reference databases facilitate species-level identification. However, these comparisons are based primarily on environmental samples, and the relative persistence of different RNA targets under digestive conditions remains poorly understood. Future studies should therefore directly compare 18S and COI targets under controlled feeding experiments.

Finally, strict separation of DNA and RNA detection remains essential. Even small amounts of DNA contamination in RNA extracts can result in false-positive RNA detection, particularly when highly abundant multicopy genes such as 18S are targeted. Robust DNase treatment and appropriate negative controls should therefore be integral components of future RNA-based dietary studies.

### Conclusions

Our study provides new insights into the feeding ecology of carabid predators in riparian systems, indicating that *Bembidion* can act as a facultative scavenger when carrion is available. We further evaluated whether RNA and DNA detected in predator gut contents can be used to trace scavenging events but found that, under our conditions, these markers did not reliably distinguish scavenging from predation. However, analysis of prey RNA and DNA offers exciting perspectives to determine feeding on live prey and may offer more reliable inference than laboratory experiments or stable isotope analysis (Keenan & DeBruyn, 2019). We outline several methodological adjustments and offer guidance for refining molecular approaches to improve the detection of carrion feeding in future studies.

## Acknowledgements

We thank Kai Riess for his help in setting up the qPCR analysis and Barbara Lösch for supporting us with fresh *D. melanogaster*.

## Author contributions

BE, FM and JS conceived the study. FM conducted the feeding experiment and AR performed molecular analysis assisted by BE. AR and BE analysed data. BE lead manuscript writing. All authors contributed to revising the final version of the manuscript.

